# Non-Canonical Binding of a Small Molecule to Sortilin Alters Cellular Trafficking of ApoB and PCSK9 in Liver Derived Cells

**DOI:** 10.1101/795658

**Authors:** Robert P. Sparks, Andres S. Arango, Zachary L. Aboff, Jermaine L. Jenkins, Wayne C. Guida, Emad Tajkhorshid, Charles E. Sparks, Janet D. Sparks, Rutilio A. Fratti

## Abstract

Sortilin regulates hepatic exocytosis and endocytosis of ApoB containing lipoproteins (ApoB-Lp) and mediates the secretion of the subtilase PCSK9. To elucidate connections between these pathways, we previously identified a small molecule (cpd984) that binds to a non-canonical site on Sortilin. In hepatic cells cpd984 augments ApoB-Lp secretion, increases cellular PCSK9 levels, and reduces LDLR expression indicative of reduced secretion of PCSK9. We have shown that insulin-induced ApoB-Lp degradation occurs through Vps34-dependent autophagy. Here we show that the specific Vps34 inhibitor PIK-III enhances ApoB-100 secretion, reducing cellular levels of PCSK9 and Sortilin resulting in reduced LDLR expression, which implicates a role for autophagy in PCSK9 secretion. Results suggest that Sortilin is central to both PCSK9 and ApoB-100 secretion. Finally, we found that cpd984 in yeast blocks CPY secretion while increasing vacuolar homotypic fusion in a Vps10-dependent manner, indicating an evolutionarily conserved mechanism required for lysosomal protease trafficking.

## INTRODUCTION

Trafficking pathways in yeast have been studied extensively as models of eukaryotic biology, which have in turn have been informative for discoveries in mammals related to human disease ^1^. Vps10 in yeast traffics the lysosomal protease CPY (carboxypeptidase) from the Golgi to the vacuole in an anteretrograde fashion and Vps10 recycles back to the Golgi via the retromer. Vps10 orthologue Sortilin in mammals traffics proteins from the Golgi where it is activated by furin mediated cleavage of its prodomain ^2^. Sortilin is responsible for trafficking its ligands between secretory pathways and lysosomal protein pathways ^3^. Vps10 and its mammalian counterpart Sortilin release cargo upon entering acidic endosomal compartments, which may be followed by dimerization of the luminal β-propeller domains of Sortilin. Subsequently, Vps10/Sortilin can be recycled back to the Golgi by the Retromer pathway, which could be mediated by dimerization ^4^. In higher eukaryotes, Sortilin is expressed in diverse cell types where it serves as a tissue specific cargo receptor. In neuronal cells, Sortilin participates in the secretion of neurotensin (NT) and other neuromodulators ^5^, while in adipose tissue, Sortilin mediates the expression of the type 4 glucose transporter Glut4 at the plasma membrane ^6^. In hepatic cells, the Sortilin receptor primarily regulates ApoB containing lipoproteins (Apo-Lp) and the protease PCSK9 (proprotein convertase subtilisin-kexin type 9). When secreted, PCSK9 binds to LDL receptor (LDLR), which targets both LDLR and PCSK9 for lysosomal degradation ^7^.

Through GWAS analysis, SORT1 mutations have been identified as candidates that correlate mutations in Sortilin with cardiovascular disease outcomes ^8,9^. Sortilin has complex relationships with lipoprotein metabolism ^10^ and in lipid accumulation in arteries ^11^. At the plasma membrane, Sortilin is found in Clathrin-coated pits similar to the LDLR, both of which bind ApoB-Lp ^12^. Differences in function and ligand specificity for the two receptors is unknown. Further complicating understanding of Sortilin function is its additional role in hepatic VLDL secretion where hepatic knockouts of Sortilin have been described to either increase or decrease VLDL secretion ^13^. Sortilin is a multiligand receptor that can interact with the various ligands found on VLDL including B100 ^14^, ApoE ^15,16^, and PI(3,4,5)P_3_ ^17^. Circulating VLDL content of PI(3,4,5)P_3_ is enriched as compared to HDL and LDL^17^. The paradox for the ability of Sortilin to both increase or decrease VLDL secretion could relate to the relative concentration and position of ligands on the VLDL surface. Presence of PI(3,4,5)P_3_ on lipoproteins relates to insulin signaling where PI(3,4,5)P_3_ is considered as a principal mediator of insulin signal transduction, which could provide a mechanism for short-term modulation of VLDL interaction with Sortilin ^18 19^.

An additional lipoprotein related ligand for Sortilin is the subtilase PCSK9, which has been shown by SPR to bind to human Sortilin with high affinity ^20^. Importantly, this study showed that neurotensin (NT), a natural substrate for Sortilin, did not inhibit PCSK9 binding, even at saturating concentrations ^21^. It has been proposed that PCSK9 is chaperoned for secretion by Sortilin. Once PCSK9 is released by sortilin extracellularly, it binds to LDLR resulting in degradation of the associated complex resulting in decreased expression of LDLR on the cell surface as LDLR bound to PCSK9 results in degradation of both PCSK9 and LDLR at the lysosome. We now show that PCSK9 secretion can be modulated by targeting Sortilin. We hypothesize that the relative secretion of Sortilin and PCSK9 can be co-regulated in hepatocytes by VLDL composition and by non-canonical binding of molecules to Site-2 of Sortilin. We suggest that this pathway is a complex and dynamic system for regulating ApoB-100 metabolism, which takes into account the balance of cellular intake (LDLR and VLDLR) and secretion of VLDL, a very low-density progenitor of LDL. We further show that the non-canonical Site-2 of Sortilin. may have evolved from primitive systems for protease trafficking in yeast involving Vps10 with similarities for carboxypeptidase (CPY) and PCSK9 trafficking ^22^.

Results presented suggest a potential alternative to current strategies targeting inhibition of the binding of PCSK9 to the LDL receptor (LDLR) ^23^. We hypothesize that using Site-2 directed Sortilin binding molecules such as cpd984, hepatic LDLR expression and VLDL secretion could be mincrease under appropriate clinical settings. Furthermore, approaches could be developed to selectively regulate involved pathways by utilizing both sites of Sortilin, as we now show the inverse results of Site-1 and Site-2 specific Sortilin molecules ^24^.

## Results

### Cpd984 and Cpd541 Bind at Different Locations of Human Sortilin

Our previous studies defined the interaction of cpd984 with Site-2 on Sortilin and its effects on NT binding and VLDL secretion ^24^. Here we tested whether cpd984 in combination with cpd541, a Site-1 directed small molecule could be used to examine the relationship between Sites-1 and -2 on Sortilin. In **Figure 1A**, we show SPR results of a titration of cpd984 concentrations binding to Sortilin. This provides strong evidence that cpd984 binds Sortilin with high affinity, having a *K*_*D*_ of 118 ± 38 nM **(Fig. 1A).** Next, we determined the binding affinity of Sortilin for cpd541, which we determined has a *K*_*D*_ of 6.9 ± 1.1 µM for Sortilin **(Fig. 1B)**. To determine whether binding Site-2 is independent of Site-1, we measured the binding of cpd984 in the presence of a nearly saturating concentration of 10 µM cpd541. We found that cpd984 bound to Sortilin with a similar affinity in the presence of 10 µM cpd541 as in the absence of cpd541 **(Fig. 1C)**. Results are consistent with our previous study showing that cpd984 bound Sortilin in the presence of C-terminal NT ^17^.

**Figure 1.**
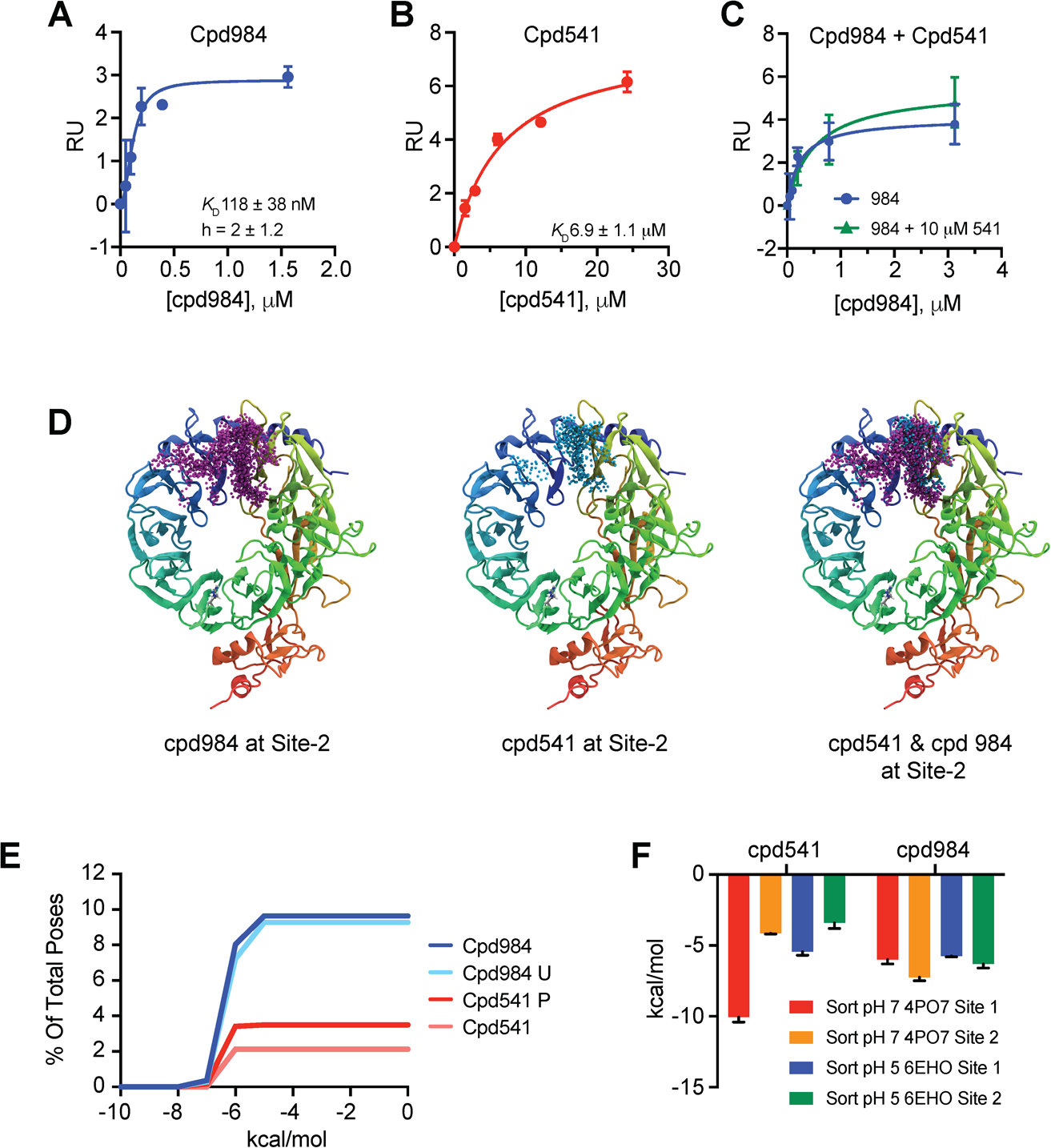
Experimental and Computational Characterization of cpd541 and cpd984. **A**, SPR determined binding affinity of cpd984 to on CM5 chip loaded with ∼3000 RU of human Sortilin with saturation curve (blue) indicating *K*_*D*_ 118 ± 38 nM. **B**, SPR as in 1a, of cpd541 to Sortilin with saturation curve (red) *K*_*D*_ 6.9 ± 1.1 µM. **C**, SPR as in 1a and 1b with cpd984 titration in the presence (green) and absence (blue) of 10 µM cpd541 with saturation curves. **D**, Resultant poses from ensemble docking for cpd984 (purple) and cpd541 (cyan) at Site-2 of Sortilin alone and superimposed. **e**, Percent of overall poses from ensemble docking versus relative docking score for cpd984 (red) and cpd541 (blue) where the Y axis is the percent of all poses for ensemble docking that bind site 2 of Sortilin, with cpd984 having about 9 % percent of all poses in site 2 and cpd541 having about 3 percent of all poses. **f**, Schrodinger Glide Xp scores of cpd541 and cpd984 docked in site 1 and site 2 of ∼pH7 PDB ID: 4PO7 and ∼pH5.5 PDB ID: 6EHO.

Using ensemble molecular docking we showed that cpd984 binds to regions of Sortilin comprising Site-2 **(Fig. 1D)**, with a higher percentage than cpd541 **(Fig. 1E)**. Furthermore, we found that the protonated carboxylate on cpd541 did not affect its preferential clustering in Site-1 versus the more hydrophobic Site-2. In parallel, we found that cpd984 with an unprotonated pentaamine ring has a basic pH of 8.4 (± 1.2), as determined by *ab initio* calculations using Jaguar ^25^. The protonation state of cpd984 had no effect on Site-2 binding, whereas cpd541 protonation state of the carboxylate resulted in a slight enrichment of cpd541 to Site 2 of Sortilin (**Fig. 1E**). To test whether pH dependent conformational changes impacted Site-1 or Site-2 binding, using Schrodinger Glide XP, we computationally docked both cpd541 and cpd984 in both Site-1 and Site-2 of a crystal structure of Sortilin at neutral pH (PDB ID: 4PO7) and acidic pH (PDB ID: 6EHO) as shown in **Figure 1F**. Cpd541 showed dramatically reduced binding to Site-1 of Sortilin as compared to cpd984, indicating that cpd541 is similar to other Site-1 substrates that bind to the NT binding site of Sortilin, and lose affinity at low pH ^26^. Furthermore, docking results confirm that cpd984 shows little discrimination between acidic and neutral pH structure of Sortilin.

Using molecular dynamics (MD), six 50 ns simulations were run for cpd541 and cpd984 for both Sites-1 and -2 comprising 24 simulations over 1 microsecond in total (**Fig. 2A**). The three best binding poses of cpd984 to Site-1 were chosen using ensemble poses generated from clusters of Site-1, whereas the three best poses with a salt bridge from cpd541 to Arg292 with the best autodock docking score were chosen and all three poses run in duplicate **Figure 1D**. For Site-2, the cpd984 pose comprising the closest match to our previously determined pose of cpd984 to Site-2 was used, whereas for cpd541 the three best poses of cpd541 to Site 2 were chosen^24^. Results and analysis of the MD simulations are presented in **Figure 2**. Cpd984 (**Fig. 2A**) and cpd541 (**Fig. 2D**) average distances from Arg292 in Site-1 and Leu539 in Site-2 are given. We found that cpd984 remained within 10 Å from Leu539 throughout the simulation (**Fig 2A)** ^24^. In contrast, simulations of cpd984 to Site-1 showed that it did not stay within 10 Å of R292 for Site-1. In addition to the stable association of cpd984 with Site-2, we found that cpd984 buries itself into the hydrophobic Sortilin beta propeller, between blades 1 and 10. Using F555 as a reference amino acid nearing the end of blade 10 of Sortilin, we show that cpd984 on average over six 50 ns simulations comes closer to this hydrophobic cavity over time (**Fig. 2B**). A representative endpoint of one of these simulations is presented in a ligand interaction diagram of cpd984 is presented in **Figure 2C**. Taken together, we hypothesize that the hydrophobic cpd984 is more likely to stay bound to Site-2 than to Site-1. Additionally, as cpd984 does not carry a negative charge at acidic or neutral pH, and has greater association with Site-2 than Site-1 of Sortilin, it is likely that cpd984 binding to Sortilin is pH independent in the cell (**Fig. 1E, F**).

**Figure 2.**
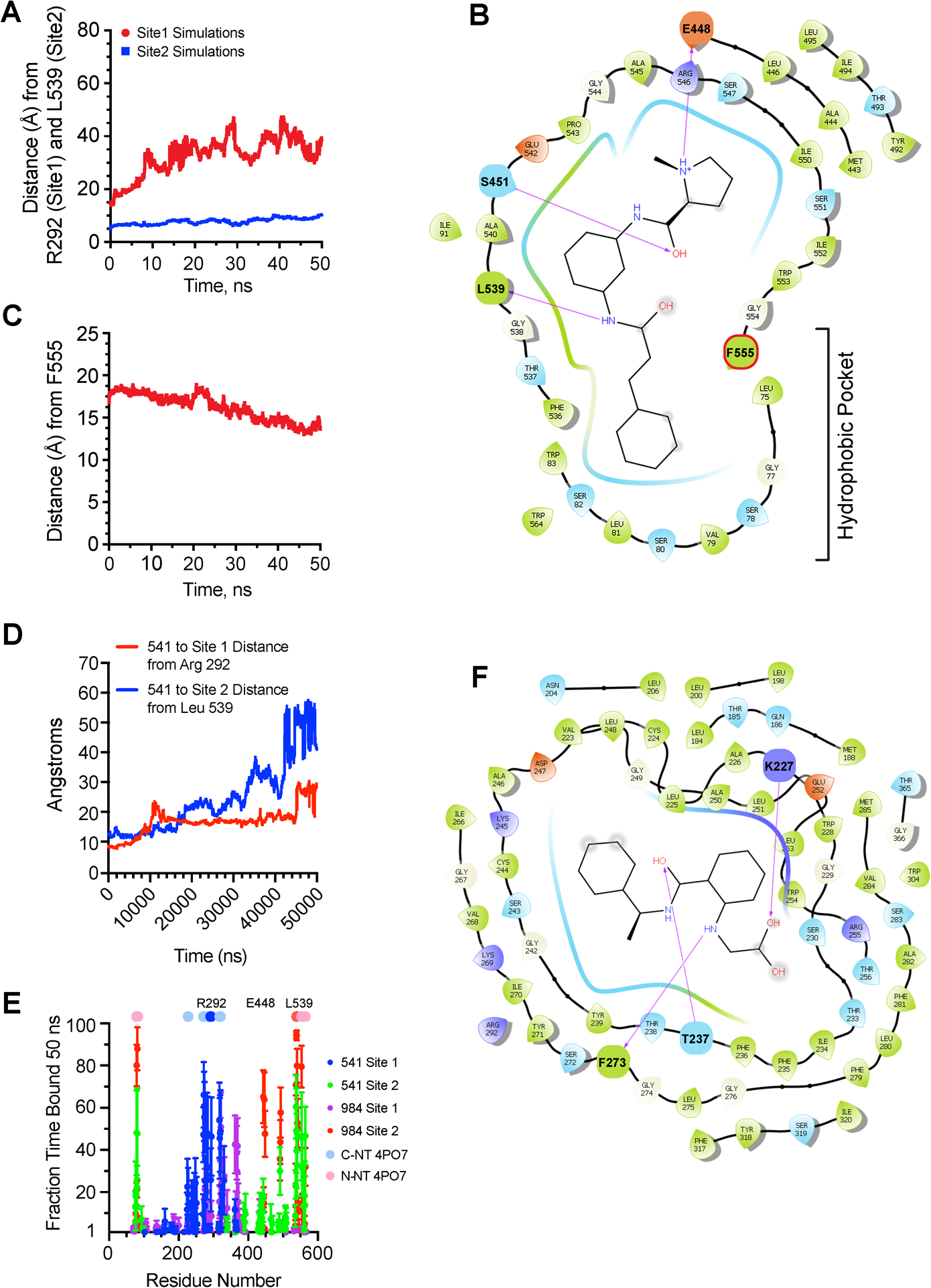
Computational Characterization of cpd984 to Sortilin Site-1 and Site-2. **A**, Simulations of bound cpd984 poses taken from ensemble docking as in Figure 1d, 1e with Site-2 pose taken from closest ensemble docking pose to pose (n=6) and the three highest scoring poses for cpd984 to Site-1 (n=2 each) based on ensemble docking. Center of mass difference over time (50 ns) between cpd984 and representative residue of Site-1 R292 for the three duplicate simulations of cpd984 to Site-1 are shown in red, whereas center of mass differences between cpd984 and representative residue of Site-2 Leu539 for the six duplicate simulations of cpd984 to Site-2 are shown in blue. **B**, Ligand interaction diagram taken from a representative simulation from Figure 2a of cpd984 to Site-2 of Sortilin at the end of the 50 ns MD simulation. **C**, Center of mass difference over time as in Figure 2a between cpd984 and a representative residue of the hydrophobic ligand binding pocket of Sortilin adjacent to Site-2 Phe555 of six simulations of Site-2 bound ligands). **D**, Simulations of bound cpd541 as in **Fig. 2A**, with center of mass taken from Arg202 and Leu539 as in **Fig. 2A**, from top 3 cpd541 ensemble docking poses in site 2 (n=2) and cpd541 poses from site 1 with a salt bridge between cpd541 and Arg292. **e**, Residue analysis for all 24 50 ns simulations of cpd984 and cpd541 in site 1 and site 2 of Sortilin with fraction time bound calculated by determining number of frames (5000) where a residue of Sortilin was within 3.5 Å of either cpd984 or cpd541. C-term and N-term NT used as a legend at 100% over the one frame taken from PDB coordinates of PDB ID: 4PO7. **F**, Ligand interaction diagram taken from a representative simulation from Figure 2d of cpd541 to Site-1 of Sortilin at the end of the 50 ns MD simulation.

Regarding cpd541, we hypothesized that it requires some electrostatic binding to form a salt bridge with Arg292 and stably interact with Site-1 via its carboxylate. In **Figure 2D**, we found that cpd541 did not stay as tightly associated to Site-2 of Sortilin as compared cpd984. In contrast, cpd541 stably associated with Site-1 more than Site-2. That said, not all cpd541 simulations remained associated with R292. A fraction of cpd541 simulations started with a salt bridge to Arg292, eventually stably forming a salt bridge with Lys227 as depicted in **Figure 2F** depicting one of the final frames of a cpd541 simulation to Site-1 of Sortilin. In **Figure 2E**, we show the fraction time bound over the course of all 24 50 ns simulations as a function of the amount of frames of the simulations as a whole, where either cpd541 or cpd984 stayed within 3.5 Å of a given residue of Sortilin, over the course of all 50,000 frames of the simulation, with each frame representing 1 ps. It is clear that cpd984 had a greater fraction bound time than cpd541 in Site-2 of Sortilin, and that cpd541 had a greater fraction time bound than cpd541 to Site-1 (**Fig. 2E**). Furthermore, the residues of interest for these interactions corresponded well to the NT peptide binding modes of Sortilin from PDB ID: 4PO7, which we have hypothesized represents the full NT peptide across the Sortilin cavity connecting Site-1 and Site-2 of Sortilin ^24^.

### Sortilin-targeting Small Molecule alters NT Binding and ApoB Secretion

Using SPR to monitor NT binding to Sortilin we found that cpd541 competed for NT binding to Sortilin in a dose dependent fashion **(Fig. 3A)**. In contrast, we now show that cpd984 enhanced NT binding to Sortilin in a dose dependent fashion ^24^. As NT binding has been shown to be mainly guided by Site-1, we hypothesize that cpd541 as an inhibitor most likely binds to NT binding Site-1 of Sortilin ^27^. Next, we examined the effects of these compounds on ApoB secretion. This was done with different cell lines in which Sortilin levels were differentially reduced with siRNA. Sortilin knockdown cell lines 1-4 expressed 95%, 70%, 40% and 10% of Sortilin, respectively, relative to the scrambled siRNA control cells **(Fig. 3B)**. As previously reported, cpd984 enhanced ApoB secretion in all cell lines, in proportion to the amount of Sortilin expressed **(Fig. 3B)**. In contrast, cpd541 reduced ApoB secretion consistent with Site-1 being the primary binding region of Sortilin for VLDL. When administered together, cpd984 and cpd541 restored VLDL secretion to levels near the untreated control **(Fig. 3C)**. This suggests that binding Site-2 allosterically regulates Site-1 binding.

**Figure 3.**
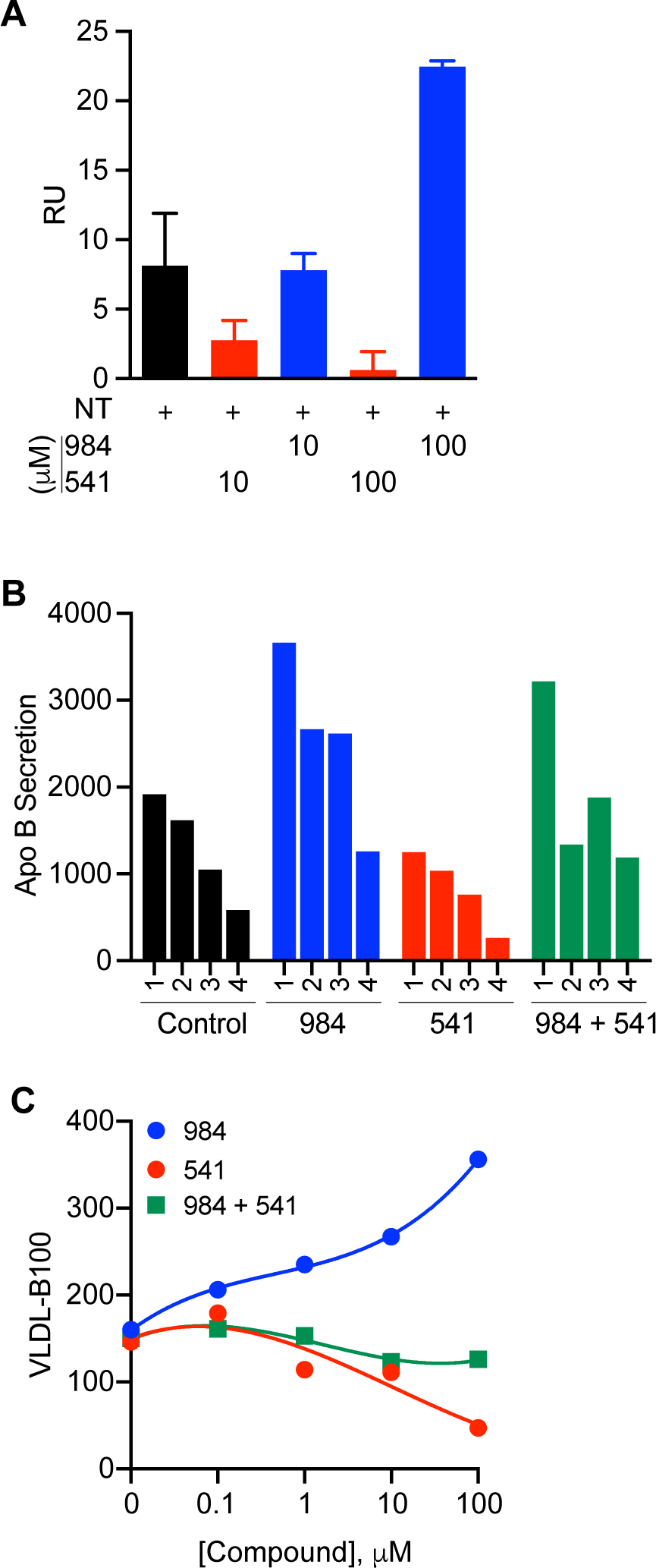
Cpd984/cpd541 Differential Effects on NT Binding and ApoB Secretion ion McA Cells. **A**, SPR analysis following administration of 100 nM NT alone and in the presence of either cpd984 or cpd541 at concentrations of 10 and 100 μM. RU subtractions of cpd984 and cpd541 alone were performed for injections at the corresponding concentrations in the presence of neurotensin in order to depict the effect of these compounds on the binding of neurotensin to Sortilin**. B**, McA knockdowns with varying levels of Sortilin were measured for secretion of VLDL-B100 into media and assessed by immuno-slot blot. The effect of Sortilin *K*_*D*_ on VLDL-B100 secretion and VLDL-ApoB secretion by insulin sensitive McA cells (1% BSA/DMEM). Results are the average of triplicate plates for each condition ± S.D. * *p*<0.05. **C**, Insulin sensitive McA were incubated with increasing concentrations of cpd541 or cpd984 and secretion of VLDL-B100 into media was assessed by immune-slot blotting.

### The Vps34-Specific Inhibitor PIK-III blocks Autophagy and Sortilin Function

Previous studies of Sortilin indicate a specialized autophagy responsible for insulin-dependent degradation of ApoB in a manner independent of p62 ^28,29^. To test effects of Vps34 specific effects on trafficking regulation of VLDL and LDLR we used the Vps34-specific inhibitor PIK-III in lieu of pan-PI 3-Kinase inhibitors such as wortmannin ^30^. In order to determine efficacy of PIKIII on autophagy as a control, we measured the autophagy markers LC3-I, LC3-II, and p62 in the presence or absence of wortmannin or PIKIII. Basal autophagic flux was blocked with E/P (E-64d and Pepstatin) ^31,32^, or HCQ (hydroxychloroquine) ^33,34^, or left untreated. Autophagy activation was detected by presence or absence of LC3-II. We found that LC3-II production was inhibited completely under all three conditions with PIK-III, whereas wortmannin had no effect **(Fig. 4A)**. Furthermore, it appears that neither PIK-III nor wortmannin affected p62 levels under autophagic conditions similar to a previous report suggesting a selective form of autophagy associated with Sortilin ^28^. As a measure of insulin-triggered autophagy, we tested whether PIK-III had an effect on the phospho-AKT pathway. We found that PIK-III, unlike wortmannin, failed to inhibit phospho-AKT production (**Fig. 4B)**.

**Figure 4.**
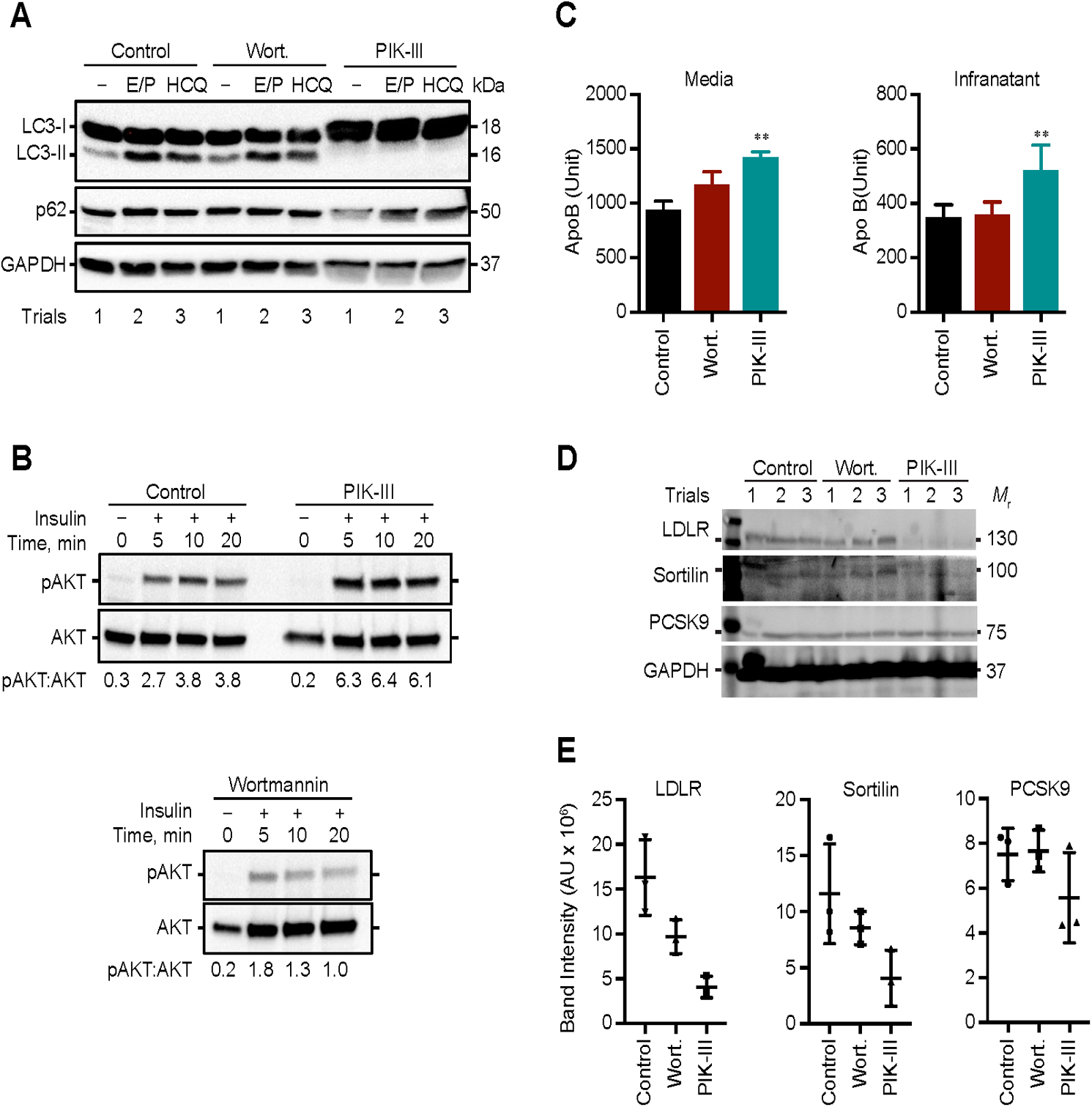
PIK-III Increases Secretion of Both ApoB and PCSK9. **A**, Measurement of autophagic protein expression in the presence and absence of wortmannin and PIK-III Cellular proteins were analyzed by IB for LC3-I, LC3-II, p62, and GAPDH using HRP-linked secondary antibody and ECL detection. Loading controls included GAPDH (cb1001mAb (6C5)). McA cells were incubated in 1% BSA/DMEM with DMSO, wortmannin or PIK-III for 18 h. Band intensities were measured using ChemiDocXRS+ system and evaluated using ImageLab 5.1 software. **B**, Measurement of AKT pathway as in Figure 4a with (5, 10, and 20) and without insulin (0) in the presence and absence of PIK-III and wortmannin blotting for both AKT and pAKT with ratios depicted on the bottom. **C**, Measurement of total ApoB as in Figure 4a in both the media and infranatant in the presence and absence of PIK-III and wortmannin. **D**, Measurement of cellular Sortilin, LDLR (LS-C146979, 1:2000), PCSK9 (Ab125251, 1:2000) and GAPDH as in Figure 4a, in the presence and absence of PIKIII and wortmannin. **E**, Quantification of LDLR, Sortilin, and PCSK9 as in 4a.

### PIK-III Alters PCSK9 Secretion and LDLR Surface Expression

After characterizing PIKIII as an autophagy inhibitor, we tested for its effects on Sortilin mediated trafficking by monitoring both VLDL secretion and PCSK9 regulation of LDLR in the presence and absence of wortmannin and PIKIII. Our results indicate that PIK-III affected both VLDL secretion (**Fig. 4C**) and LDLR expression (**Fig. 4D**). It appears from **Figure 4C** that total ApoB increased with PIKIII in both the cell and the media. We hypothesize that as PIKIII inhibits autophagic degradation of ApoB, it would then increase the total amount of ApoB available, therefore, we would expect more ApoB shifted towards secretion while the intracellular increase in ApoB was due to lack of autophagic degradation. Immunoblotting for intracellular pools of PCSK9, Sortilin, and LDLR showed a decrease in LDLR expression, and reduced levels of Sortilin and PCSK9 (**Fig. 4D**). We hypothesize that decreased intracellular PCSK9 is consistent with increased PCSK9 secretion, resulting in decreased LDLR expression. The decreased intracellular Sortilin is likely due to the requirement for more Sortilin to be diverted to secretion due to increased ApoB availability.

### Effect of cpd541 and cpd984 on the Sortilin-VLDL-PCSK9 Trafficking Pathway

We further tested the effect of modulating Sortilin trafficking through defined Site-1 and Site-2 molecules. We found that the Site-2 cpd984 enhanced LDLR expression **(Fig. 5A**), whereas cpd541 had little effect on PCSK9. We hypothesized that PCSK9 binds to Site-2, which could explain why cpd984 was effective in modulating PCSK9 as opposed to cpd541. Using the protein-protein docking prediction software ClusPro, we docked PDB ID: 4PO7 Sortilin to PDB ID: 2P4E PCSK9, removing the NT fragments of 4PO7 prior to simulation. All 10 clusters showed PCSK9 binding to Blade-1 of Sortilin (**Fig. 5B**), and selected the top 3 poses with the lowest overall predicted ΔG. We ran the three generated clusters for 25 ns each and analyzed both Sortilin and PCSK9 contact points for binding approximated by having a distance of less than 3.5 Angstroms between the two proteins. This showed that the residues of Sortilin closest to PCSK9 over throughout multiple trajectories remained to be adjacent to Site-2 or on the outside of the Sortilin cavity opposite Site-2. In **Figure 5D**, we show a root-mean-squared fluctuation (RMSF) analysis of these same trajectories for Sortilin. Residue analysis (**Fig. 5C**), and RMSF analysis (**Fig. 5E**) were additionally performed for PCSK9 using residues at its interface with LDLR for reference utilizing a 5 ns MD simulation of PCSK9 to LDLR from PDB ID: 3P5C. All results are consistent with PCSK9 binding to Site-2.

**Figure 5.**
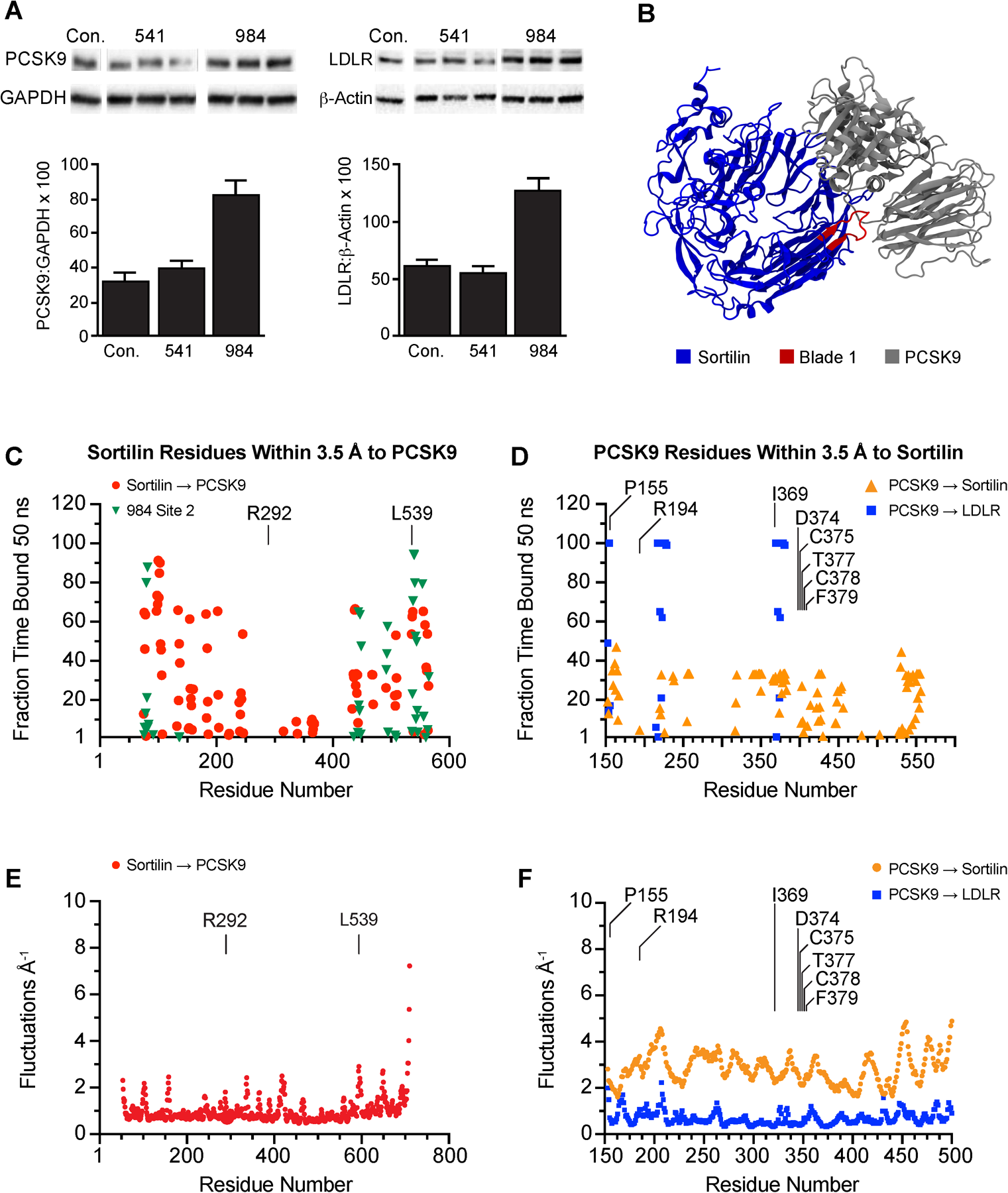
Effects of Cpd541 and Cpd984 on PCSK9/LDLR Expression and Potential Role of PI(3,4,5)P_3_ in Sortilin Binding to ApoB-Lp. **A**, McA cells were incubated in 1% BSA/DMEM with DMSO, cpd541 (10 μM) or cpd984 (10 μM) for 18 h. Cellular proteins were extracted and analyzed by IB for LDLR (LS-C146979, 1:2000) or PCSK9 (Ab125251, 1:2000) using HRP-linked secondary antibody and ECL detection. Loading controls included – β-Actin (Rockland ph600-401-886) and GAPDH (cb1001mAb (6C5)). Band intensities were measured using ChemiDocXRS+ system and evaluated using ImageLab 5.1 software. **B**, Results from ClusPro protein-protein docking simulation of Sortilin PDB ID: 4PO7 to PCSK9 PDB ID: 2P4E with Sortilin in blue, PCSK9 in grey, blade 1 of Sortilin in red. **C**, Residue analysis as in Figure 2e corresponding to three 10 ns simulations of PCSK9 to Sortilin using PDB files in Figure 5b with mean fraction bound as in Figure 2e for PCSK9 to Sortilin with 984 results from Figure 2e plotted in green, PCSK9 residue. **D**, RMSF analysis of Sortilin residues from ClusPro simulation of Sortilin PDB ID: 4PO7 to PCSK9 PDB ID: 2P4E e, RMSF Analysis of PCSK9 residues from ClusPro simulation in Figure 5E.

### Characterization of VLDL and LDL Sortilin Affinity

Having concluded previously that PI(3,4,5)P_3_ binds to Site-1 ^17^, we tested *in vitro* binding using SPR to determine the effect of cpd541 and cpd984 on Sortilin binding of PI(3,4,5)P_3_. First, we found that diC8-PI(3,4,5)P_3_ bound with high affinity to crosslinked Sortilin with a *K*_*D*_ of 4.2 µM ± 380 nM (**Fig. 6A**). When we ran the same diC8-PI(3,4,5)P_3_ concentration curves in the presence of 10 µM cpd541 and 10 µM cpd984, we found that cpd541 abolished PI(3,4,5)P_3_ binding, whereas cpd984 enhanced diC8-PI(3,4,5)P_3_ binding about 10-fold to Sortilin with a *K*_*D*_ of 474 Nm ± 85 nM. To determine if these effects held true for more native forms of PI(3,4,5)P_3_, we used nanodiscs containing 2.5% PI(3,4,5)P_3_. Sortilin bound nanodiscs with an increased binding affinity over diC8-PI(3,4,5)P_3_ for Sortilin with a *K*_*D*_ of 55 nM ± 13 nM indicating importance for the lipid membrane to Sortilin binding. Similarly to diC8-PI(3,4,5)P_3_, the affinity of this interaction was enhanced about 10-fold by cpd984 to a *K*_*D*_ of 5.4 nM ± 0.8 nM **(Fig. 6B)**.

**Figure 6.**
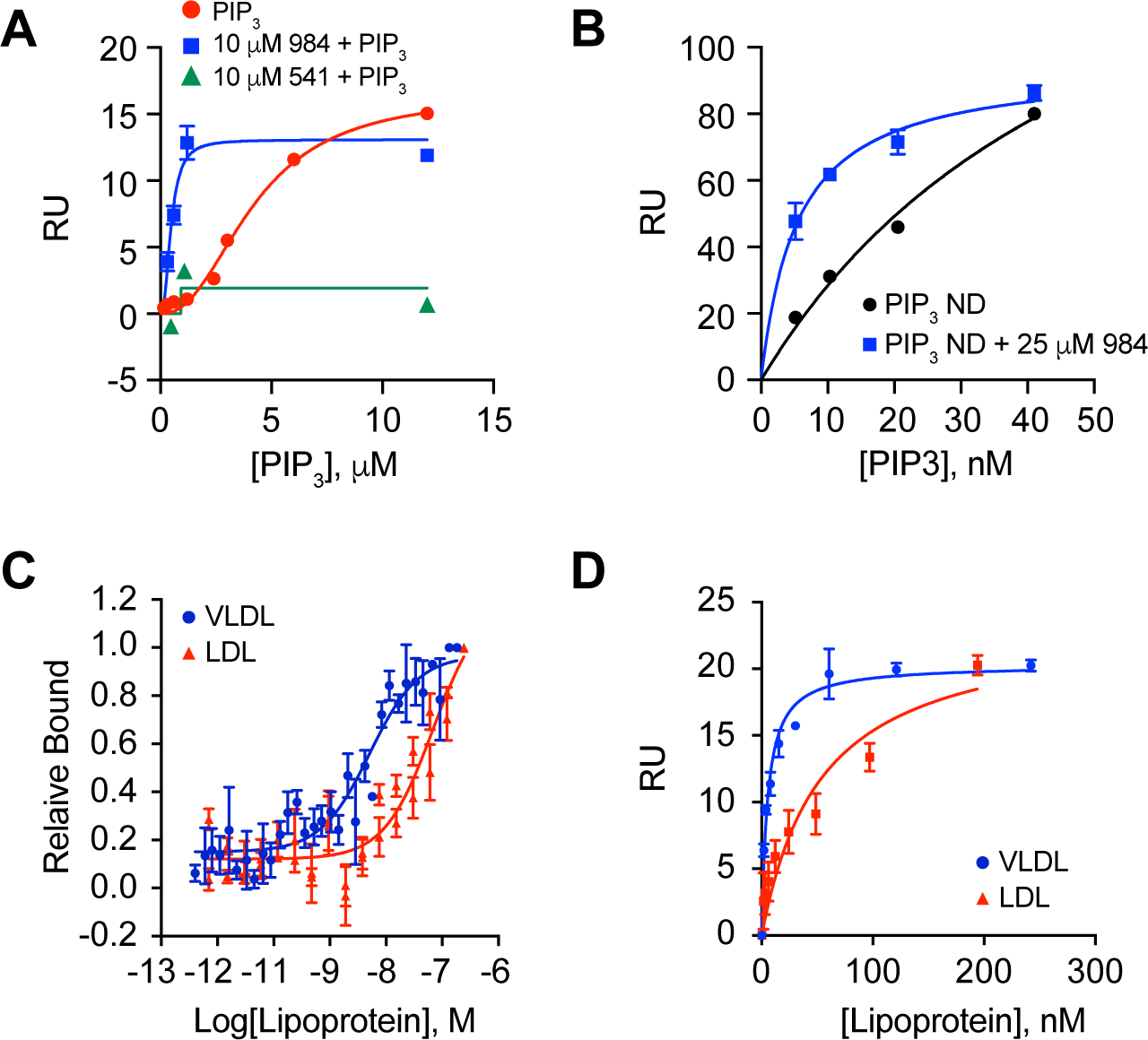
Cpd984 modulates binding to PI(3,4,5)P_3_. **A**, SPR as in Figure 1a of diC8-PI(3,4,5)P_3_ binding measurements in the presence and absence (red) of 10 µM cpd984 (blue) and cpd541 (green) with *K*_*D*_. **B**, SPR using a CM5 chip crosslinked with Sortilin. **C**, Labeled microscale thermophoresis (MST) using Atta-647 Ni-NTA labeled Sortilin measured in the red channel with a constant concentration of Sortilin of 12.5 nM and 32 different concentrations of VLDL and LDL run in triplicate and analyzed using Nanotemper M.O. Affinity Analysis software with *K*_*D*_ for VLDL of ∼4 nM and for LDL of ∼74 nM. **D**, SPR performed using a CM5 chip containing ∼2000 RU of Sortilin with *K*_*D*_ for VLDL of ∼5 nM and for LDL of ∼54 nM.

We previously showed that circulating VLDL contains more PI(3,4,5)P_3_ than LDL ^17^. Using these same fractions, we characterized rat derived circulating VLDL and LDL binding to Sortilin using SPR and MST. Here we show that rat VLDL bound Sortilin with a *K*_*D*_ of ∼4 nM, an order of magnitude higher relative to binding LDL with a *K*_*D*_ of 54-74 nM. We hypothesize that this was due to differences in particle composition, where VLDL acquired PI(3,4,5)P_3_ during the co-synthesis with ApoB. (**Fig. 6D, E**). These data suggest the differences between LDL and VLDL binding was independent of ApoB, as both particle types contain the protein, suggesting a primary difference was the presence of PI(3,4,5)P_3_.

### Sortilin Small Molecule Effect in Yeast

The yeast gene *VPS10* encodes the type I transmembrane receptor for carboxypeptidase Y (CPY) that cycles between the late Golgi and the prevacuolar compartment ^22,35^. The CPY hydrolase enters the ER as an inactive zymogen and is modified by four N-linked polysaccharides forming p1CPY (67 kDa) ^36^. Thereafter, p1CPY transits through the Golgi where polysaccharides are extended to form the p2CPY precursor (69 kDa). p2CPY is transported from late Golgi in vesicles containing the subtilase Kex2, and is subsequently delivered to the vacuole. Upon arrival, the amino pro-segment of p2CPY is cleaved to yield enzymatically active mature CPY or mCPY (61 kDa). Similar to Sortilin, Vps10 has been shown to have multiple binding sites for different ligands in yeast, however, unlike Sortilin, Vps10 contains two protein sorting domains making Vps10 substantially larger than human Sortilin. ^37^

We hypothesized that Sortilin trafficking of PCSK9 should share consistencies with Vps10 sorting of CPY, reasoning that the mechanism for regulation of trafficking of proteases destined for the lysosome or vacuole could help elucidate Sortilin cellular itinerary for other ligands such as VLDL-ApoB. First, we deleted *VPS10* from our fusion tester yeast strains DKY6281 and BJ3505 and compared vacuolar levels of CPY and the late endosomal/lysosomal rab GTPase Ypt7. In **Figure 7A** we show immunoblots of isolated vacuoles from *vps10*Δ yeast and their parent strains. These results indicate that CPY at the vacuole was reduced in the absence of Vps10 in both strains.

**Figure 7.**
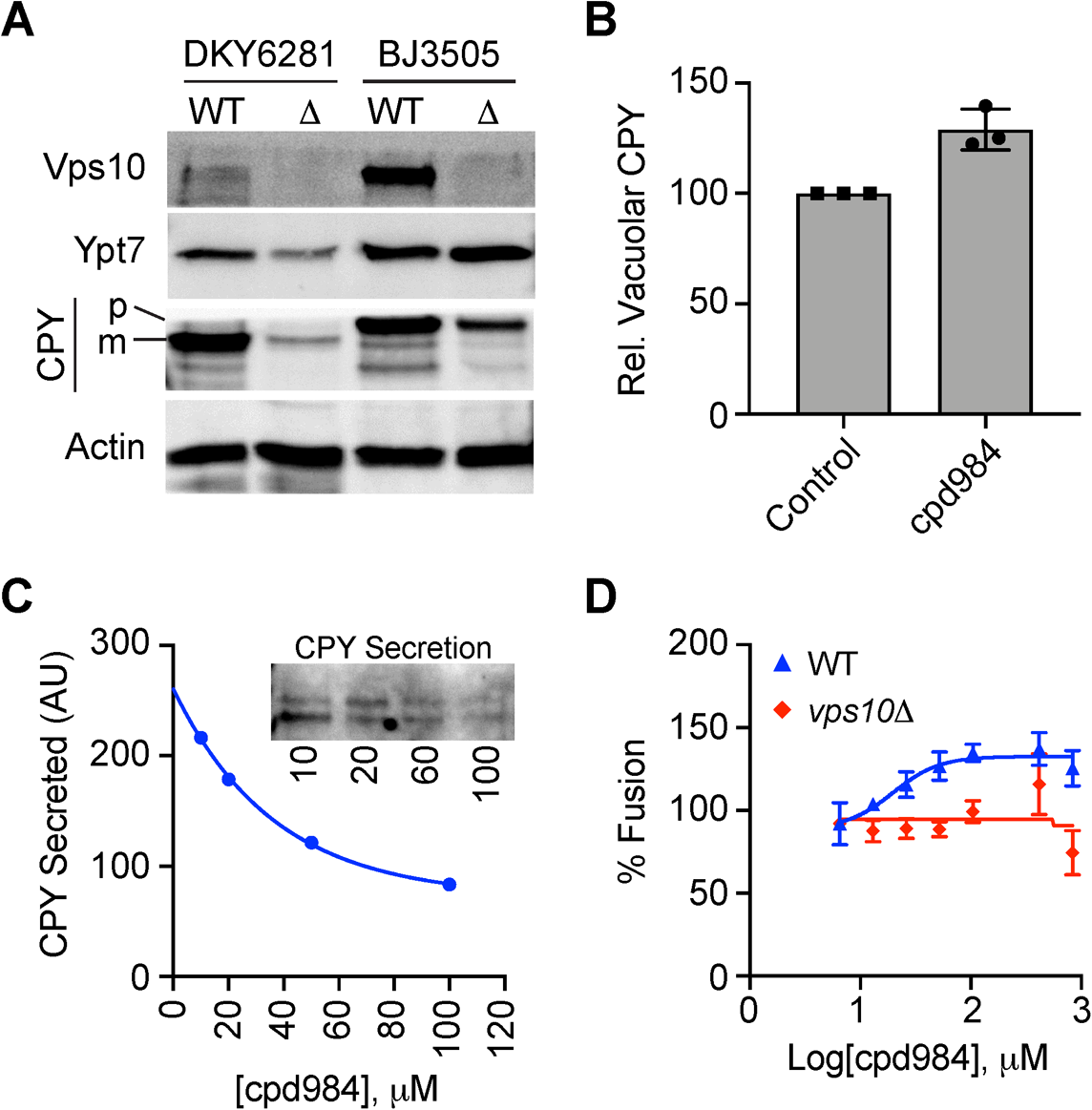
cpd984 affects CPY sorting in Yeast. **A**, Immunoblots of wild type and *vps10*Δ cell lysates with antibody against the late endosomal/lysosomal Rab Ypt7, the yeast Sortilin ortholog Vps10, the luminal protease CPY, and actin as a loading control**. B**, Relative concentrations of intracellular CPY in untreated WT cells vs cells treated with cpd984. **C**, Immunoblot of secreted CPY in cells treated with cpd984. **D**, In vitro homotypic vacuole fusion assays with wild type and *vps10*Δ vacuoles in the presence or absence of cpd984.

Like PCSK9, we assume that increases in CPY at the vacuole, correspond with decreases in secretion of CPY. We used a classical technique to characterize trafficking defects in response to cpd984 ^38^. To test if a Site-2 homologous area was present on Vps10, we treated wild type yeast cells with cpd984 and measured CPY secretion. We found that cells treated with 20 µM cpd984 accumulated substantially more CPY at the yeast vacuole relative to the untreated control **(Fig. 7B)**. To test whether the increase in vacuolar CPY correlated to a decrease in secreted CPY stores, we treated individual yeast cultures with a concentration curve of cpd984. As shown in **Figure 7C**, secreted CPY was reduced with increased cpd984 concentrations. Together these data showed that a Site-2-like domain may be present on Vps10.

Finally, we hypothesized that altered trafficking to the vacuole via Vps10 increases in vacuolar CPY due to cpd984 could alter vacuolar homotypic fusion as a measure for increased trafficking to lysosomes. In vitro vacuole fusion assays showed that cpd984 administration resulted in dose dependent increases in fusion of WT vacuoles, while no effect was seen with *vps10*Δ vacuoles, suggesting that proteins responsible for changes in fusogenicity may depend on Vps10 (**Fig. 7D**). This lends support to a hypothesis that cpd984 enhances Vps34 recruitment resulting in increased trafficking to the lysosome as a potential mechanism.

## Discussion

The endocytic pathway for ApoB-Lp uptake by LDLR in hepatocytes has been studied extensively and forms much of the basis for current therapies controlling high LDL cholesterol in humans ^39^. Little is known about the role of Sortilin in this process or how the relative concentrations of these receptors are regulated by Sortilin in this pathway ^7,10^. In this study we undertook to further examine the role of Sortilin in LDL/VLDL trafficking. These data show that Sortilin binds VLDL with higher affinity compared to LDL (**Fig. 6C, D**). We speculate this to be the result of increased ligands for Sortilin on the VLDL surface other than ApoB-100, e.g. PI(3,4,5)P_3_ or ApoE, in contrast to decreased alternative ligands for LDL other than ApoB-100 ^17,40^.

Considering that PCSK9 is an additional ligand for Sortilin-mediated secretion in hepatocytes ^7^, we further explored the effect of cpd984 on PCSK9 binding to Sortilin. Using functional endpoints in McA cells we demonstrated that cpd984 treated cells retained PCSK9 (**Fig. 5A)** resulting in increased expression of LDLR. The absence of these effects with the Site-1 ligand cpd541 suggests Site-2 mediates this function. These results are consistent with the hypothesis that Site-2 is the primary region of Sortilin for PCSK9 in chaperoning its secretion. Furthermore, we showed that VLDL secretion is inversely modulated by administration of these two compounds (**Fig. 3C**), that both cpd effects depend on Sortilin expression levels (**Fig. 3B**), and that when both sites are occupied effects on VLDL secretion are attenuated (**Fig. 3C**). This indicates the potential for site specific modulation of sortilin chaperone activity.

Regarding NT binding to Sortilin, we showed that cpd541 inhibits NT binding whereas cpd984 enhances binding (**Fig. 3A**). Results suggest a potential allosteric effect of Site-2 binding on VLDL binding to Site-1, resulting in increased secretion (**Fig. 3B, C**). Sortilin plays a central role in hepatic ApoB-Lp metabolism, yet its exact role remains enigmatic ^13^. The questions that remain are: 1) How do Sortilin knockdowns both increase and decrease hepatic VLDL secretion? ^10^; 2) Do separate VLDL ligands interact with one another when binding to Sortilin?; and 3) How important is Sortilin in mediating ApoB-Lp endocytosis through clathrin coated vesicles? Considering the complexity of these questions, we propose a model where Sortilin contains at least two interactive sites for ligand binding.

In addition to mediating ApoB-Lp secretion, Sortilin has a significant role in ApoB degradation by the lysosome ^41^. Our previous work showed that blocking autophagy prevents insulin signaling from suppressing ApoB secretion. Whether insulin dependent autophagy is merely an accumulation of basal autophagy is unknown, however insulin-dependent PI(3,4,5)P_3_ production its incorporation into VLDL leads us to posit that the production of this lipid diverts ApoB-Lp to the autophagic pathway. We further demonstrated that autophagy regulates PCSK9, Sortilin and LDLR levels (**Fig. 4C, D**). This process is not dependent on the production of pAKT triggered by insulin, as PIK-III failed to block AKT phosphorylation (**Fig. 4B**) ^30^. The reduction in LDLR, Sortilin and PCSK9 in the presence of PIK-III suggests that loss of autophagy results in increased secretion of Sortilin and PCSK9 with resultant PCSK9-mediated reduction in LDLR. Results suggest loss of autophagy affects Sortilin related processes involving both Site-2 and Site-1.

Considering the importance of Site-2 in hepatic ApoB-Lp metabolism, we speculated that the Site-2 related pathways for PCSK9 and ApoB-Lp may be conserved in other cell types with some probability that it could have evolved from a primitive pathway present in yeast. The mechanistic role of the yeast CPY protease receptor Vps10, the ancestor of Sortilin, is well well-documented, and also involves at least two ligand binding domains ^42^. We used cpd984 to ask whether it might modulate CPY trafficking as it had with Sortilin with respect to the protease PCSK9. Indeed, treating yeast with cpd984 resulted in decreased CPY secretion while increasing its intracellular pool, suggesting CPY sorting is a parallel pathway to PCSK9 sorting as shown by its accumulation at the vacuole and decrease extracellularly (**Fig. 7B, C**). It appears that these effects are Vps10 dependent as fusion increases with cpd984 were attenuated in *vps10*Δ strains (**Fig. 7D**).

These studies demonstrate a complex interconnected regulatory system for hepatic lipoprotein metabolism based on evolution of Vps10. Considerable research will be required to completely define this mode of regulation. Considering the complexity of this system, we believe that our observations on Sortilin regulation in hepatocytes and yeast may serve as a paradigm for Sortilin trafficking in other cell types such as neuronal and adipose. Implications of these studies may also have impact on understanding other Sortilin related processes including insulin secretion ^8,24,43^, Glut4 transport ^44–47^, and plaque formation in Alzheimer’s ^48–50^ as well as serving as a model for function of other Vps10 homologues such as Vth1, Vth2, and YN94 or Sortilin homologues such as SORCS and SORLA involved in similar processes as Sortilin. Understanding and delineating these protein functions should improve our understanding of the complex regulation of proteins involved in Vps10 family sorting pathways ^51,52^

## Methods

### Cell culture, materials and reagents

McArdle RH-7777 cells (McA cells) were cultured as previously described in serum containing complete Dulbecco’s Modified Eagle’s Medium (cDMEM) ^43,53.^ To induce insulin sensitivity, McA cells at 50-60% confluency were incubated for 18 h in serum-free media consisting of DMEM containing 1% (w/v) BSA (1% BSA/DMEM). Human Sortilin (Sortilin) (Ser78-Asn755) with C-terminal 6-His tag was from R&D Systems, Inc., (Minneapolis, MN). Plasma from fasted Sprague Dawley rats (BioreclamationIVT, Westbury, NY) was used to prepare VLDL ApoB standards. Mouse monoclonal antibody to rat B100 was prepared in our laboratory and characterized previously ^54^. Rabbit anti-Sortilin antibody was from GeneTex (GTX54854, Irvine, CA), BD Medical (A311709, Franklin Lakes, NJ) or from Abcam (ab16640, Cambridge, MA). Rabbit polyclonal PCSK9 and LDLR antibody was from LSBio (Seattle, WA). Mouse anti-glyceraldehyde phosphate dehydrogenase (GAPDH) from Santa Cruz Biotechnology. Rabbit anti-p62/SQSTM1 was from (Medical & Biological Laboratories, Nagoya, Japan) and anti-LC3 from (MBL International, Woburn, MA). Lipofectamine 2000, Plus™ Reagent, and rabbit anti-Vps34 antibody were from Invitrogen/ThermoFisher. Anti-rabbit and anti-mouse horseradish peroxidase (HRP)-linked IgG and Hyperfilm™ were purchased from GE Healthcare. Rabbit anti-p-AKT (9271) and rabbit anti-AKT (9272) were from Cell Signaling Technology (Danvers, MA). Antibody against yeast Ypt7 and actin were previously described ^55^, and CPY (10A5B5, ThermoFisher). Mouse anti-pY Platinum 4G10 was from EMD Millipore (Temecula, CA). Horseradish peroxidase linked donkey anti-rabbit IgG (NA9340), sheep anti-mouse IgG (NXA931) and ECL Prime Western Blotting Detection Reagent (RPN2232) were from GE Healthcare. All other materials and reagents were essentially as described previously ^53^. diC8-PI(3,4,5)P_3_ (1,2-dioctanoyl-phosphatidylinositol 3,4,5-triphosphate) was from Echelon Inc. Wortmannin was from Sigma Chemical Corp and 4’-(cyclopropylmethyl)-N2-4-pyridinyl-[4,5’-bipyrimidine]-2,2’-diamine (PIK-III) from Cayman Chemical (Ann Arbor, MI). Compound 98477898 (2S)-1-methyl-N-3-[(3-phenypropanoyl)-amino]phenylpyrrolidine-2-carboxamied (cpd984) and compound 54122218 [2-([(1R)-1-phenylethyl]aminocarbonyl)phenyl]amino}acetic acid were obtained from ChemBridge Corp. (San Diego, CA). Stock solutions of cpd984 and cpd541 (10 mM) were prepared in DMSO, and stored in aliquots at −20°C.

### Cell Culture

Rat hepatocytes (RH) were isolated from Sprague-Dawley rat livers, and were cultured on collagen-coated dishes in Waymouth’s 751/1 medium containing 0.2% (w/v) BSA as described previously ^56^. Wild-type McA cells were maintained in culture in complete DMEM (cDMEM) ^53^. Inhibitors were used at reported concentrations for cpd984 and cpd541. PIK-III was administered at 1 µM and wortmannin administered 10 µM. Inhibitors were validated in RH where cell toxicity was minimal as determined by LDH release.

### Knockdown of Sortilin in McA cells using siRNA

McA cells were transfected using Fugene6 according to manufacturer’s protocol (Promega Corp., Madison WI) using three different pGIPZ based vectors expressing shRNAi targeting rat Sort1 mRNA (V2LMM_58553, V3LMM_450660, V3LMM_450662), and one scrambled, non-silencing control (GE Healthcare Dharmacon, Lafayette, CO) as previously described ^24^. Puromycin selection was performed on McA cells. Sortilin knockdown (KD) from each cell line was examined by immunoblotting.

### Immunoblotting

McA cell lysates were prepared and denatured proteins were separated by SDS-PAGE as described previously ^43^. Membranes were incubated with primary antibodies overnight at 4°C in blocking buffer with antibody binding detected with species specific secondary HRP-linked antibodies and developed using Amersham™ Prime reagent (GE Healthcare). Insulin signaling to AKT evaluated with Bio-Rad nitrocellulose membranes with phosphospecific (pY, p-AKT(Ser473) and mass specific antibodies ^28^. Chemiluminescence was measured with ChemiDocXRS+ system (Bio-Rad) and quantified using Image Lab 3.0.1 software from Bio-Rad (Hercules, CA). Autophagy was measured under three conditions: 1. HPQ (hydroxychloroquine) leads to the accumulation of basal LC3-II by blocking fusion between autophagosomes and lysosomes ^33,34^; 2. E/P (E-64d and Pepstatin-A) prevents the fusion of autophagosomes the lysosomes ^32^; and 3. The control were wild type McA.

### Immuno-slot blotting

VLDL and LDL preparations for SPR and MST in Figure 5 were prepared as previously described ^24^Experimental media were adjusted to 1% (v/v) protease inhibitor cocktail I (EMD Millipore) and to a salt density of 1.019 g/ml by addition of a solution of NaBr (d = 1.495 g/ml). VLDL was isolated by ultracentrifugation in a L-70 Ultracentrifuge (Beckman Coulter, Inc., Fullterton, CA) using a 50Ti rotor (200,000 x g, 18 h, 14°C). Following centrifugation, the top 1.5 – 2.0 ml VLDL fraction was removed using a syringe and weighed to determine volume. VLDL samples were applied in triplicate wells (0.2 to 0.4 ml per well) in a Bio-Dot® SF apparatus (Bio-Rad). Two PVDF membranes were used together for blotting: the top was Immobilon-P (IPVH09120 SF) and the bottom was Immobilon-PSQ (ISEQ09120 SF); both were obtained from EMD Millipore. VLDL-ApoB standards were prepared from rat plasma VLDL and total ApoB (B100 and B48) and B100 content were determined on stained gels following SDS-PAGE separation ^57^. VLDL-ApoB standards in TBS were slotted in duplicate alongside test samples. After filtration, 0.4 ml of TBS was added as a wash. After final filtration, membranes were air dried, rehydrated in methanol and incubated in TBS then in blocking buffer at 4°C overnight. At this stage slot blots were evaluated similarly to immunoblotting. After chemiluminescence development and B100 quantitation, slot blots were stripped by incubation in Restore™ PLUS (ThermoFisher) for 15 min at room temperature and were reblocked overnight at 4°C. Total ApoB (B100 and B48) present in VLDL was then evaluated following incubation with rabbit polyclonal anti-rat ApoB. The bottom PVDF membrane was carried through the entire procedure to assure there was no “bleed through” of test samples. VLDL-ApoB and VLDL-B100 content were calculated from standard curves generated by VLDL-ApoB standards. Recovery of rat VLDL added to unspent media averaged 94% ± 4.9% with a CV of 5.2% (n = 6 replicates).

### Computational modeling and compound screening

Schrödinger’s Maestro program (version 9.3.5) was used as the primary graphical user interface and Maestro version 10.2 (Schrödinger, LLC, New York, NY) was used for ligand interaction diagramming. Virtual screening was performed on compounds contained in ChemBridge libraries (www.chembridge.com) that were prepared with Schrödinger’s LigPrep program (Schrödinger, LLC, New York, NY). The virtual screening method was performed using Schrödinger’s GLIDE software ^58^ on the Sortilin crystal structure PDB ID: 4PO7 27. Compounds were docked on grids generated with Glide with cpd541 docked at a box determined by C-terminal NT and cpd984 docked at a box determined by the N-terminal fragment of NT. Grids were then adapted from alignment of PDB ID:6EHO to PDB ID: 4PO7 and docking performed for all grids using Glide XP setting with results exported into Graphpad Prism.

### MD Simulations of apo and holo Sortilin

Using the aforementioned crystal structure of Sortilin (PDB ID:4PO7) molecular dynamics simulations were done using NAMD 2.12 ^59^ using the CHARMM36m force field ^60^. Prior simulating, the system was prepared using the CHARMM-GUI solution builder, with a salt concentration of 150 mM NaCl. Simulation parameters included constant pressure of 1 atm via Langevin dynamics, as well as a constant temperature of 310 K using Langevin piston Nosé-Hoover methods ^61,62^. Long-range electrostatic forces were evaluated using the particle mesh Ewald (PME) with a 1 Å grid spacing ^63,64^. Van der Waals interactions were calculated using a 12 Å cutoff with a force-based switching scheme after 10 Å, as well as a 2 fs time step integration via the SETTLE algorithm ^65^. Visualization and analysis was done using VMD 1.9.3 ^66^. The system was equilibrated for 20 ns restraining the Cα atoms of the protein (1.0 kcal/mol/Å^2^) to allow for solvation. This was followed by a production run of 50 ns without restraints for 4 poses taken from ensemble docking, 3 poses with the highest affinity pose for Site-1 of Sortilin run in duplicate for each, and the pose for Site-2 that represented the predicted pose as shown previously ^17^.

### Ensemble Molecular Docking of cpd541 and cpd984

To probe cpd541 and cpd984 interactions with Sortilin, ensemble molecular docking was employed as described ^67^. Using snapshots from the 50 ns production simulation to sample protein dynamics, snapshots were taken every 200 ps. Each of the resultant 250 snapshots were used to dock cpd541 and cpd984 using a 100Å by 90Å by 70Å grid box. Docking was done with an exhaustiveness of 10, yielding a total of 2500 docked poses. Resultant poses were clustered using a hybrid K-centers/K-medoids algorithm, utilizing an RMSD method ^68,69^. Representative poses with highest scoring affinities in clusters closest to Site-1 and Site-2 were selected for further 50 ns simulations. The resultant drug bound simulations were analyzed with VMD as well as MDAnalysis ^70,71^.

### Protein-Protein Docking and Simulations

Characterization of protein-protein interactions between Sortilin (PDB:4PO7) and PCSK9 (PDB:2P4E) was done using the default settings of webserver ClusPro ^71-73^. The top three output poses where simulated for a production run of 10 ns using MD protocols as for Apo and Holo Sortilin. Interacting residue analysis was done using VMD with a 3.5 Å cutoff, over each of the 10 ns trajectories and exported into GraphPad Prism for analysis. Root mean squared fluctuations (RMSF) of Cα atoms of the proteins Sortilin and PCSK9 were analyzed and exported from VMD and plotted using GraphPad Prism.

### Microscale Thermophoresis

Thermophoresis measurements were performed using a Monolith NT.115 labeled thermophoresis instrument ^72,73^. Sortilin-His6 was labeled with Ni-NTA Atto-488 according to the manufacturer’s protocol as previously described ^73,74^. M.O. Control software was used for operation of MST. Target protein concentrations were 50 nM for Sortilin labeled protein. LED excitation power was set to 90% and MST set to high allowing 3 sec prior to MST on to check for initial fluorescence differences, 25 sec for thermophoresis, and 3 sec for regeneration after MST off. Analysis was performed using M.O. Affinity Analysis Software as the difference between initial fluorescence measure in the first 5 sec as compared with thermophoresis at 15 sec. All measurements were performed in PBS buffer (137 mM NaCl, 2.7 mM KCl, 8 mM Na_2_HPO_4_, and 2 mM KH_2_PO_4_, pH 7.4) without Tween and binding affinity was generated using Graphpad Sigmoidal 4PL fit from points exported from M.O. Affinity Analysis software using *K*_*D*_ Model with target concentration fixed at 50 nM.

### Surface Plasmon Resonance

Surface plasmon resonance measurements were performed on a Biacore T200 instrument equipped with CM5 sensor chips with ∼2000 response units (RU) of Sortilin covalently immobilized to the surface for VLDL and LDL binding, ∼3500 RU crosslinked Sortilin for small molecule binding, and a CM5 with ∼6500 RU crosslinked Sortilin for both NT and nanodisc binding. Lipid composition of nanodiscs consisted of 3.023 µmol dipalmitoyl phosphatidylcholine (PC), 0.098 µmol diC16-PIP3, and 0.78 µmol 1-palmitoyl, 2-oleoyl phosphatidylethanolamine (PE), which were prepared as previously described ^17^. HBS-DMSO running buffer (10 mM HEPES pH 7.4, 150 mM NaCl, 1% DMSO) was used at a flow rate of 30 μl/min and injections performed with times for association of 90 sec, and dissociation of 300 sec, followed by injection of buffer to regenerate the Sortilin surface. Regeneration for Ni-NTA non-covalently linked nanodisc experiments utilized EDTA ^24^. Regeneration for CM5 NHS/EDC crosslinked Sortilin required a 30 sec injection of 10 mM NaOH as previously described ^17^. Binding was expressed in relative RU; the difference in response between the immobilized protein flow cell and the corresponding control flow cell. NT, cpd984, 5% PI(3,4,5)P_3_ containing nanodiscs and VLDL/LDL were administered to chips containing Sortilin with 1:1 titrations and results exported from BiaEvaluate software into GraphPad prism (GraphPad Software). Cpd984, cpd541 and PI(3,4,5)P_3_ saturation curves were fit using a specific binding equation with hill slope, whereas all other SPR saturation curves were fit using a 1:1 specific binding model.

### Vacuole Isolation and in vitro vacuole fusion assay

Vacuoles were isolated from BJ3505 ^75^ and DKY6281 ^76^, which were used for fusion assays by density gradient floatation as previously described ^72^. Fusion reactions (30 µL) contained 3 µg each of vacuoles from BJ3505 (*pep4*Δ *PHO8*) and DK6281 (*PEP4 pho8*Δ), fusion assay buffer (125 mM KCl, 5 mM MgCl2, 20 mM PIPES-KOH pH 6.8, 200 mM sorbitol), ATP regenerating system (1 mM ATP, 29 mM creatine phosphate, 0.1 mg/ml creatine kinase), 10 µM CoA, and 283 nM Pbi2p and buffer or cpd984. Reactions with or without cpd984 were incubated at 27°C for 90 min and the Pho8 activity was measured in 250 mM Tris-Cl PH 8.5, 0.4% Triton X-100, 10 mM MgCl2, 1 mM p-nitrophenyl phosphate. Fusion-dependent alkaline phosphatase maturation was measured by the amount of p-nitrophenylate produced. p-Nitrophenylate absorbance was measured at 400 nm.

### Yeast Western Analysis and CPY Secretion

Vacuoles of BJ3505, DK6281, and their *vps10*Δ derivatives were isolated and vacuoles were removed by centrifugation (16,000 × *g*, 5 min, 4°C) and SDS sample buffer was added to the supernatants. Samples were heated at 95°C for 5 min, resolved by SDS-PAGE, transferred to nitrocellulose, and probed by Western blot in. CPY secretion was measured with vacuoles from BJ3505 and SEY6210 yeast strains incubated with cpd984 were grown at a volume of 1L overnight as done for yeast fusion. A 10 mL sample of YPD media was taken before harvest and spun down to remove cell lysate. TCA precipitation was performed resuspending pellet in approximately 100 µL SDS sample buffer, run on SDS-PAGE and blotted for CPY ^77^. The remaining 990 mL of YPD was used for harvesting vacuoles. Vacuoles were measured using Bradford reagent and normalized prior to running SDS-PAGE along with a loading control.

### Statistics

Unless noted, results are expressed as the mean ± S.E.M., where n equals the number of independent experiments in which replicate analyses were performed in each experiment. Significant differences were assessed using Student’s t-test with p-values ≤ 0.05 being considered significant.

## Acknowledgements

This paper is dedicated to Janet DeHoff Sparks the wife of Charles Edward Sparks and mother of Robert Pleasants Sparks. Following training at the University of Pennsylvania and Post-Doctoral training at Wistar, Janet went on to become a Full Professor at University of Rochester Medical Center in the Department of Pathology. Janet loved to get her work done early in the morning, go to Midtown Athletic Center, and then go right back to work to perform experiments. Janet continued this routine until her death in Sarasota, Fl from an unexpected heart attack that may have been due to an as yet well characterized genetic lipoprotein disorder. Many of the experiments in this paper were performed by Janet herself, not a graduate student. Furthermore, many of the experiments were also performed by Janet in the presence of her son Robert, who hung on to her proverbial coat tails attempting to get access to the latest data while performing minimal manual labor.

This research was supported by grants from the National Institutes of Health (R01-GM101132 to RAF, and P41-GM104601, U01-GM111251 and U54-GM087519 to ET), the National Science Foundation (MCB 1818310 to RAF), and the Office of Naval Research (ONR N00014-16-1-2535 to E.T.). Computational resources were provided by XSEDE (XSEDE MCA06N060) and Blue Waters (ACI-1440026). SPR was aided by the help of Dr. Jermaine Jenkins at the University of Rochester Structural Biology & Biophysics Facility with support from NIH NCRR grant 1S10 RR027241, as well as NIH NIAID P30AI078498 and the University of Rochester School of Medicine and Dentistry.

## Author Contributions

RPS, JDS, ASA, CES and RAF conceived the project and designed experiments. RPS, JDS, ASA, ZLA, and JLK performed the experiments and analyzed data. JDS, CES and RAF supervised the research. RPS, ASA, JDS, CES and RAF wrote the manuscript with input from all authors.

## Data availability

All data generated or analyzed during this study are included in this published article.

## The abbreviations used are

ApoB-Lp: ApoB containing lipoproteins
PI(3,4,5)P_3_: phosphatidylinositol (3,4,5)-trisphosphate
Sortilin: human Sortilin
NT: neurotensin
CtermNT: C-terminal neurotensin
NtermNTC: N-terminal neurotensin modified from carboxyl to amide C-terminal
MST: microscale thermophoresis
VLDL: very low density lipoprotein
B100: apolipoprotein B derived from unedited *Apob* mRNA
SPR: surface plasmon resonance
CPY: carboxypeptidase Y
McA: McCardle cells

